# Human Fibroblast-Myeloid cell tissue atlas across lung, synovium, skin and heart

**DOI:** 10.1101/2025.03.04.641204

**Authors:** Lucy MacDonald, Olympia Hardy, Melpomeni Toitou, Gabriela Kania, Przemyslaw Blyszczuk, Raphael Micheroli, Thomas D Otto, Mariola Kurowska-Stolarska, Caroline Ospelt

## Abstract

Single-cell RNA sequencing (scRNAseq) of human tissues has expanded our understanding of the complexity of cellular subsets and their changes in disease. The availability of scRNAseq data in different tissues and disease states provides an opportunity to compare cellular subsets and identify common and unique cellular activation. In this study, we aimed to characterize shared and tissue-specific myeloid and stromal phenotypes and uncover key cellular subtypes involved in pathogenic tissue activation. We analyzed scRNAseq data from 14 public datasets, comprising heart, lung, skin, and synovium in both healthy and diseased states. Our analysis identified distinct and overlapping myeloid and stromal cell populations in these tissues. Despite significant inter-individual variability, we were able to identify both shared and disease-specific changes in these cell populations. These findings provide insights into the conserved and tissue-specific roles of myeloid and stromal cells in health and disease and contribute to a better understanding of tissue pathology and potential therapeutic targets.

## Introduction

Single cell transcriptomics (scRNAseq) has begun to unravel the true complexity of cell biology by identification of novel cellular subsets through unbiased clustering analysis. The multitude of available scRNAseq data from different tissues and clinical states presents opportunities for comparison of cellular subsets across different tissues and diseases to identify common and unique transcriptional signatures. Application of integrative scRNAseq analysis recently allowed us to successfully compare macrophage populations shared between severe COVID19 and rheumatoid arthritis (RA) (1). This type of analysis is important for facilitating the repurposing of therapeutics for inflammatory diseases with multiple comorbidities.

Tissue-resident macrophages (TRMs) are critical for normal tissue development, physiology, and homeostasis (2). The majority of TRMs derive from embryonic precursors and are established in tissue before birth, where they are primarily responsible for efferocytosis of apoptotic cells and tissue debris (3, 4). However, distinct TRMs also have unique tissue-specific physiological functions (5). For instance, alveolar macrophages (FABP4+) perform steady state removal of excess lung surfactant produced by local epithelium and their deficiency results in pulmonary proteinosis (6–8). The ability of TRM to execute such crucial, unique functions is determined by signals in their local niche, which is formed by other tissue resident cells such as fibroblasts and/or epithelial cells as well as soluble mediators (9).

The phenotype of tissue resident cells is strongly influenced by loss of tissue integrity, e.g. in chronic inflammation (10). Inflammation leads to the infiltration and maturation of pro-inflammatory monocyte-derived macrophages and expansion of specific fibroblast subtypes (11, 12). Tissue-specific interaction between tissue resident macrophages and stromal cells are not well characterised up to now. Therefore, the aim of this study was to identify common and tissue-unique myeloid and stromal phenotypes (clusters) and to decipher unique and cross-tissue cells and signals involved in pathogenic tissue activation.

## Results

### Optimization of integration strategy

We collected scRNAseq data from 14 public datasets, spanning four distinct tissues (heart (13–15), lung (7, 16, 17), skin (18–20), and synovium (21–24)) including 162 healthy donors and patients (Figure 1A). We integrated datasets of each tissue using Seurat’s anchor integration (Figure 1B) and then focused our cross-tissue analysis to myeloid and stromal cell clusters. We first randomly subsampled each tissue atlas, to allow for the integration of normalized number of myeloid (n=4500), stromal (n=4500) and T cells (n=1000) from each tissue. This facilitated the identification of shared and unique macrophage and fibroblast clusters without creating a bias towards the phenotypes with the most cells. Inclusion of T lymphocytes, which are predominantly blood-derived and thus are more likely to be similar across tissues, were used to aid cross-tissue integration. As expected, UMAP visualization before integration (Supplemental Figure 1A) clearly showed that T cells (blue) from all tissues grouped as one population with considerable overlap in phenotype (Supplemental Figure 1Aii). Myeloid cells (red) from distinct tissues still grouped together (Supplemental Figure 1Ai), but with no overlap in their phenotype (Supplemental Figure 1Aii). Stromal cells (green) were most dissimilar between tissues.

**Figure 1.**
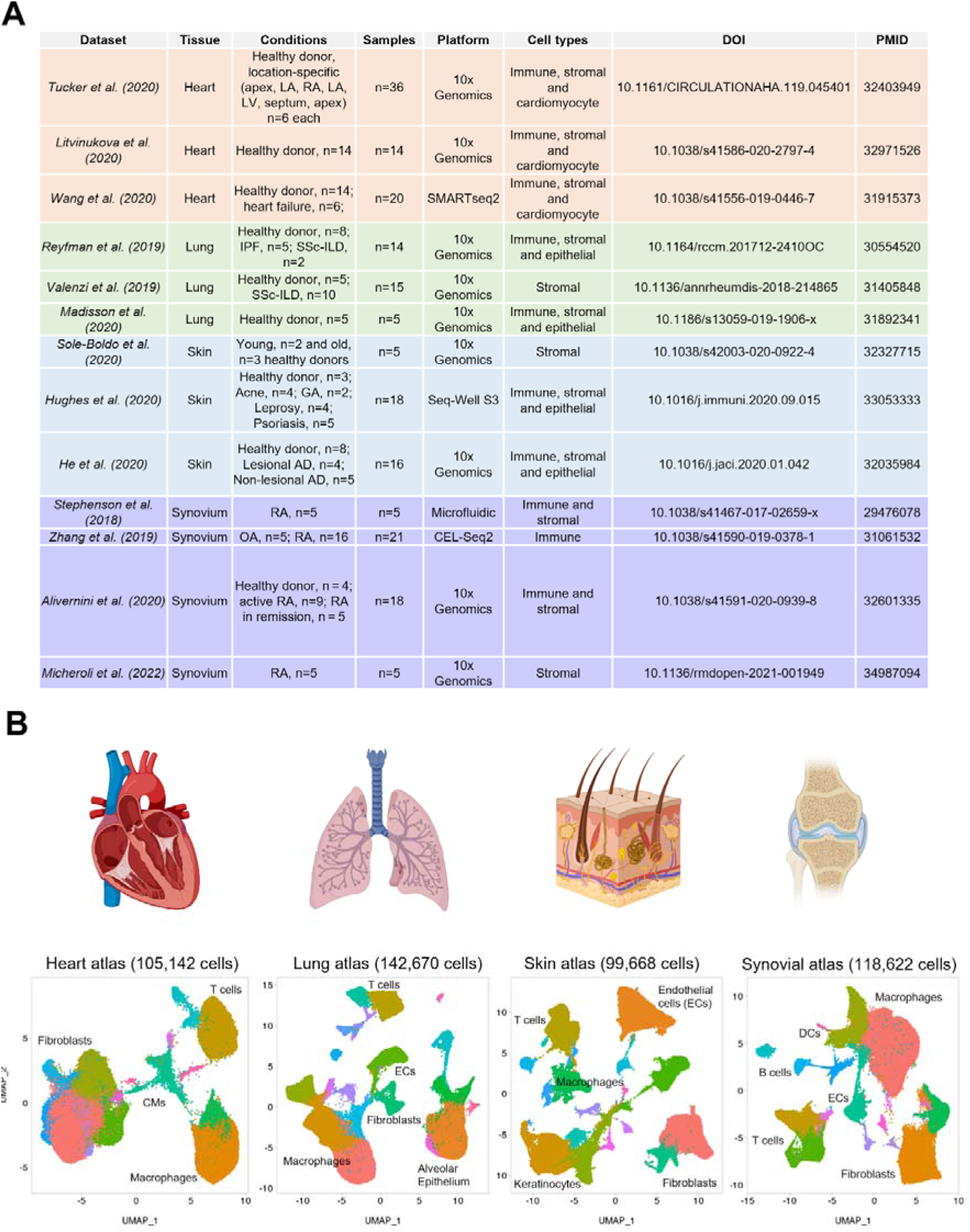
Datasets for cross-tissue integration of myeloid and stromal cells from distinct tissues/inflammatory diseases. **(A)** Table of details of datasets used in cross-tissue integration. LA=left atrium; RA=right atrium; LV=left ventricle; RV=right ventricle. **(B)** Individual UMAP visualization for the integration of datasets from heart, lung, skin, and synovium. Each cell is colored by cluster identity and prominent cell lineages are annotated.

To determine whether common myeloid or stromal cells exist across tissues, we tested three distinct approaches for removal of batch effect and compared their output, namely Seurat (25), Harmony (26) and BBKNN (27). Firstly, we applied Seurat V3 anchor integration strategy, which functions to identify pairs of similar cells across datasets (‘anchors’) and uses these pairs to correct for technical artefacts. Whilst Seurat allowed for the integration of cells from different tissues, it failed to preserve any distinctions between tissues (Supplemental Figure 1Bii). Harmony preserved biological variability whilst integrating cells from different tissues (Supplemental Figure 1Cii). BBKNN allows efficient analysis of large cell datasets by batch-correcting each cell’s k-nearest neighbours to generate an adjusted UMAP.

However, this method failed to preserve any global distances between cell lineages (Supplemental Figure 1Di) and resulted in nearly double the number of clusters (k=27) as Seurat (k=14), despite keeping the same parameters (Supplemental Figure 1B-Diii). Thus, we decided to proceed with Harmony for further analyses.

### Identification of shared and tissue-unique clusters

Having selected Harmony as our method of choice, we removed T cell populations and re-visualized the remaining myeloid and stromal cell data (Figure 2A). We identified 10 myeloid and 8 stromal cell populations across synovium, skin, lung, and heart (Figure 2B). The range of myeloid cluster included monocyte-like precursor populations (CD14+S100A12+ and CD16+ISG15+), proinflammatory (IL1B+ and SPP1+) and resolving (TREM2+, NR4A1+ and LYVE1+) macrophages as well as DC phenotypes (CCR7+, CD1c+ and CD207+) (Figure 2C). Stromal cell clusters included lubricin expressing (PRG4+) fibroblasts (Figure 2E), which were the most distinct from all other stromal cell clusters in the cluster dendrogram (Figure 2F). Nevertheless, this subtype shared some overlapping marker gene expression with the VCAM1+ fibroblast subtype (Figure 2E). Additionally, we characterised a population of CDH19+ fibroblasts, which also expressed transcripts associated with complement activation (*C7, CFD*) as marker genes and A2M+ fibroblasts that expressed *TCF21*, an important factor in lung morphogenesis(28). Other stromal clusters that we identified included collagen-producing (SPARC+), and pro-inflammatory (APOE+) fibroblasts, producing *CXCL12* and *C3*, as well as a *CD34*+ fibroblast population (MFAP5+) (Figure 2E). Finally, we found *actin* (*ACTA2*), *transgelin* (*TAGLN*) and *myosin light chain* (*MYL9*) expressing myofibroblasts-like cells (ACTA2+).

**Figure 2.**
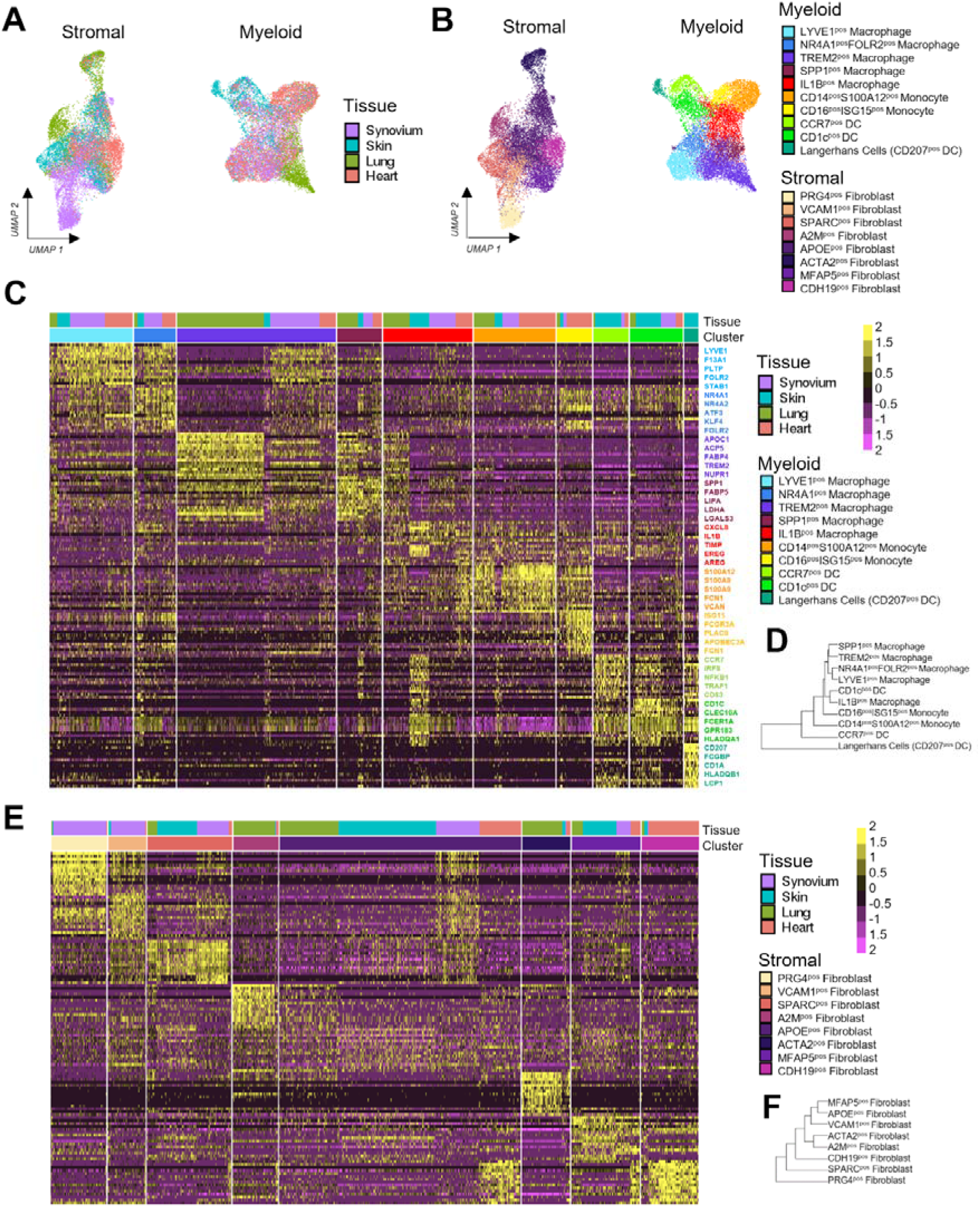
Normalized Harmony integration of heart, lung, skin and synovial tissue myeloid and stromal cells. **(A-B)** UMAP visualization of 20,000 myeloid and 20,000 stromal cells from cross-tissue integration with Harmony. Cells are represented by individual dots and coloured by **(A)** tissue origin and **(B)** cluster identity. **(C)** Heatmap of the top 15 differentially expressed genes (DEGs) per myeloid cluster. DEGs identified using MAST, with markers expressed in at least 40% of cells in that cluster. Genes are considered significantly DE if the adjusted P < 0.05 by Bonferroni correction and multiple test correction (multiplied by number of tests). Top 5 cluster markers are annotated. **(D)** Dendrogram visualization of hierarchical clustering of myeloid cell populations. **(E)** Heatmap of the top 15 differentially expressed genes (DEGs) per stromal cluster. **(F)** Dendrogram visualization of hierarchical clustering of stromal cell populations.

We then investigated how these myeloid and stromal cell populations were distributed between the distinct tissues. Analysis of myeloid cell clusters (Figure 3A-D) indicated that the heart had a high proportion of CD14+S100A12+ and CD16+ISG15+ monocyte-like precursors. The lung myeloid cell compartment was governed by resolving TREM2+ macrophages. Alternatively, the skin myeloid atlas mostly comprised DC phenotypes including CCR7+ and CD1c+ populations, and langerin-expressing (CD207+) DC, also known as Langerhans cells, which were exclusive to the skin (Figure 3B and C). In synovium, we found mixed myeloid fractions without tissue specific dominance of one cell cluster. This myeloid cell distribution was also evident when we visualized the abundance of each identified myeloid cell population throughout the analysed tissues (Figure 3D).

**Figure 3.**
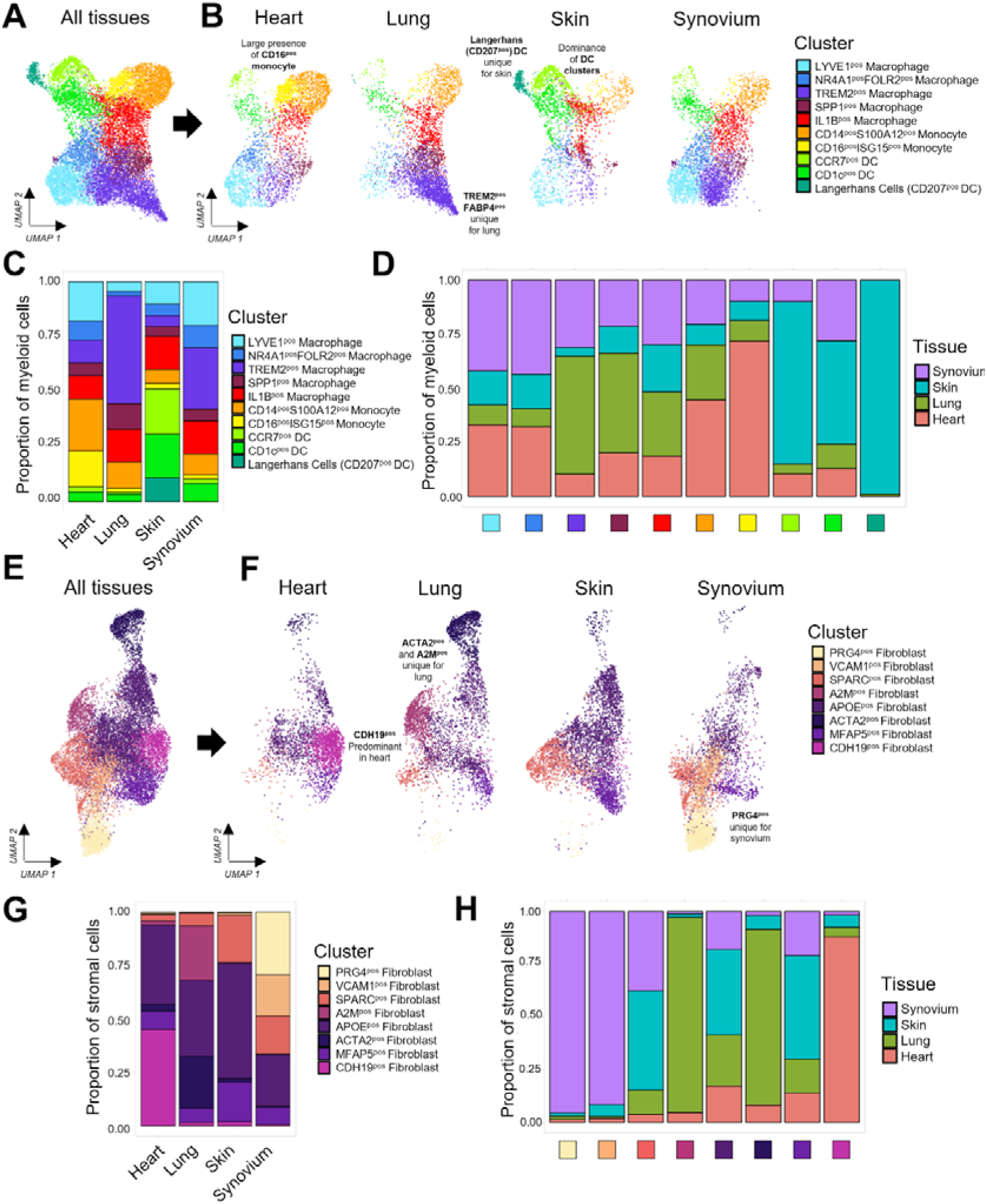
Common and unique heart, lung, skin and synovial tissue myeloid and stromal cells clusters. **(A)** UMAP visualization of 20,000 myeloid cells from cross-tissue integration with Harmony. Cells coloured by cluster identity. **(B)** UMAP visualization split by tissue of origin. **(C)** Stacked bar plot of relative proportion of cells in each myeloid cluster across tissue atlases. **(D)** Stacked bar plot of relative proportion of cells from different tissue that constitute each myeloid cluster. **(E)** UMAP visualization of 20,000 stromal cells from cross-tissue integration with Harmony. Cells coloured by cluster identity. **(F)** UMAP visualization split by tissue of origin. **(G)** Stacked bar plot of relative proportion of cells in each stromal cluster across tissue atlases. **(H)** Stacked bar plot of relative proportion of cells from different tissue that constitute each stromal cluster.

Langerhans (CD207+) DC were exclusively found in the skin and also the other DC phenotypes were derived largely from skin samples. LYVE1+ and NR4A1+ macrophages were mostly from the synovium and heart, whilst TREM2+ and SPP1+ phenotypes mainly originated from the lung. The heart was the major source of CD16+ISG15+ monocytes. Analysis of the relative proportion of stromal cell clusters across tissues (Figure 3 E-H) revealed that the heart stromal compartment was predominantly composed of CDH19+ and APOE+ fibroblasts (Figure 3G) and that CDH19+ fibroblasts were almost exclusively found in the heart (Figure 3H). In the lung stromal atlas, mainly A2M+, APOE+ and ACTA2+, but also SPARC+ and MFAP5+ populations were found (Figure 3G). A2M+ and ACTA2+ fibroblasts were lung specific (Figure 3H). The skin and synovial tissue shared the collagen-expressing SPARC+ fibroblasts as well as a proportion of APOE+ and MFAP5+ populations. No skin specific fibroblast population was found. Finally, the synovium contained the PRG4+ and VCAM1+ populations (Figure 3G), which were exclusive to this tissue (Figure 3H). These two subtypes resemble those previously described as synovial lining fibroblasts (24). In summary, our analysis identified populations of macrophages and fibroblasts that were common to multiple tissues whilst also identifying tissue-unique clusters. PRG4+/VCAM1+ synovial fibroblasts, ACTA2+ and A2M+ lung fibroblasts and CDH19+ heart fibroblasts were tissue-specific fibroblasts and Langerhans (CD207+) DC in the skin tissue specific myeloid cells. In general, tissue-specificity was more pronounced in the stromal than in the myeloid compartment.

### Changes in cell population induced by disease

We next aimed to investigate how the proportions of common and tissue unique myeloid and stromal cell clusters change between healthy donors and disease patients. To perform a comprehensive analysis of cluster proportions across disease states with sufficient numbers of cells per sample, we applied the Harmony method to integrate the total number of T lymphocytes, myeloid (136,741 cells), and stromal cells (117,102 cells) from our tissues of interest, with no limit on cell number. Again, the T lymphocyte population that served as an additional anchor was removed after integration. In total, we performed combined analysis of 253,843 myeloid and stromal cells from heart, lung, skin, and synovium (Supplemental Figure 2A). We conducted unbiased clustering and identified the clusters based on our previous nomenclature. Additional emerging clusters were characterised by marker gene expression (Supplemental Figure 2B and Figure 2E).

Integration of total myeloid cells across tissue atlases (136,741 myeloid cells) revealed an additional population of ISG15+ macrophages, which we characterised previously in the synovium (21). The total integration resulted in a change in relative proportion of some of the clusters (Supplemental Figure 2C and D). We observed an increase in the relative proportion of cells in the LYVE1+ and TREM2+ clusters and a decrease in the proportion of cells in CCR7+ DC, CD207+ DC and CD16+ISG15+ clusters (Supplemental Figure 2C).

Nevertheless, the skin was still the tissue with the highest proportion of DC populations. The proportion of cells in the LYVE1+ cluster in the heart strongly increased, while the proportion of CD14+S100A12+ and CD16+ISG15+ monocytes decreased in the heart (Supplemental Figure 2D).

Our new analysis of stromal cell clusters identified three new clusters, CARMIN+, SPSB1+ and CALD1+ fibroblasts, respectively (Supplemental Figure 2E). The relative proportion of cells in each cluster showed a substantial increase in the size of the CDH19+ cluster, mainly in the heart, and a decrease in the size of APOE+, SPARC+ and PRG4+ clusters after integration of the total cell number (Supplemental Figure 2F). The total integration also revealed that the VCAM1+ cluster was much smaller than previously anticipated (Supplemental Figure 2G). The proportion of the CDH19+ phenotype increased in all tissues, but still dominated the heart tissue (Supplemental Figure 2G). The newly emerging CARMIN+ and SPSB1+ fibroblasts were very small cell populations almost exclusively appearing in the heart. The CALD1+ population was shared between lung, skin and synovium (Supplemental Figure 2G).Thus, integration of all cells from the tissue atlas, revealed further heterogeneity of myeloid and in particular stromal cell populations.

We then went on to analyse whether and how the proportion of these identified stromal and myeloid cell populations were affected by disease. We assessed cell proportions in heart failure (HF; n=6), idiopathic pulmonary fibrosis (IPF; n=5), systemic sclerosis-associated interstitial lung disease (SSc; n=12), acne (n=4), leprosy (n=4), psoriasis (n=5), granuloma annulare (GA; n=2), atopic dermatitis (AD; n=4), osteoarthritis (OA; n=5), rheumatoid arthritis (RA; n=35) and RA in sustained clinical remission (n=5).

Concerning myeloid cell populations, LYVE1+ macrophages represented the dominant myeloid phenotype in the heart as compared to other healthy tissues (Figure 4A). In the skin, this population appeared to be slightly increased in AD, but this was statistically non-significant (Figure 4A). The NR4A1+ population (Figure 4B), the closest relative of LYVE1+ on the dendrogram (Figure 2D), was most prominent in the healthy synovium compared with skin and was minimally present in lung and heart. Its proportions tended to increase in OA as well as in acne. Similarly, the other resolving phenotype, the TREM2+ macrophage was significantly higher in the healthy synovium as compared to other tissues, but alternatively, was also highly abundant in lung tissue (Figure 4C). The ISG15+ macrophage cluster (Figure 4D), which was only identified with the updated integration of all cells, was highest in lung but appeared in other tissues with disease (AD, leprosy (skin) and OA, RA (synovium). However, its proportion remained <2% of the total myeloid population. The pro-inflammatory SPP1+ macrophage population (Figure 4E) showed the most prominent changes in healthy versus diseased tissues. It was highest in the healthy lung, yet it represented <10%. Its phenotype increased significantly with SSc-ILD. In addition, whilst this population was not present in healthy synovium, it significantly increased in active RA. Alternatively, the IL1B+ pro-inflammatory phenotype (Figure 4F) was highest in skin in health but was significantly reduced in AD and its proportion did not increase significantly in any of the analysed diseases.

**Figure 4.**
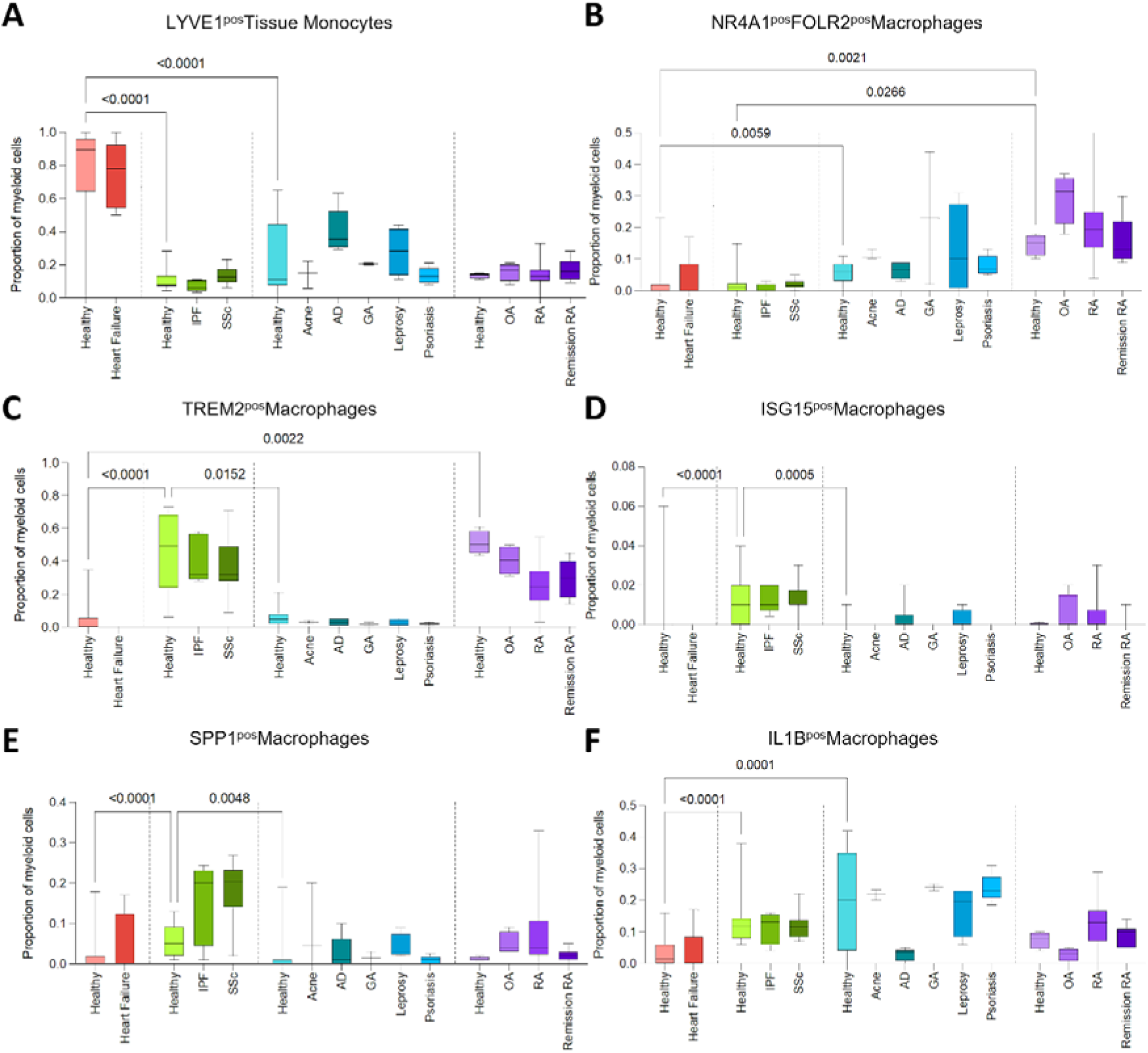
Distinct macrophage cell clusters dominate different tissues and disease states. Box and whisker plots comparing the relative proportion of cells in each myeloid cluster across different tissues healthy donors and disease states. Statistical analysis applied ANOVA with Kruskal-Wallis test for multiple comparison to compare healthy state of each tissue. Significant P values (*P* < 0.05) are shown. Heart failure (HF; n=6), idiopathic pulmonary fibrosis (IPF; n=5), systemic sclerosis-associated interstitial lung disease (SSc; n=12), acne (n=4), leprosy (n=4), psoriasis (n=5), granuloma annulare (GA; n=2), atopic dermatitis (AD; n=4), osteoarthritis (OA; n=5), rheumatoid arthritis (RA; n=35) and RA in sustained clinical remission (n=5). Red = heart; green = lung; blue = skin; purple = synovium.

The precursor CD14+S100A12+ tissue monocyte population was present in all tissues in health but at low levels (∼10%)(Supplemental Figure 3A). It was significantly increased in healthy lung versus heart but was not statistically significant in any of the disease states. The other monocyte population, CD16+ISG15+ monocytes, was also significantly higher in healthy lung compared to heart Supplemental Figure 3B). However, in some healthy heart donors and patients in remission of RA this proportion was >10% of the myeloid cells. As for the DC phenotypes, all clusters were mainly found in the skin compared to other tissues (Supplemental Figures 3C-E). The CCR7+ phenotype was significantly increased in psoriasis patients (Supplemental Figure 3C). In AD, the proportion of CD1c+ DC was significantly decreased (Supplemental Figure 3D), while the Langerhans cell population (CD207+) showed an increased proportion in AD patients (Supplemental Figure 3E).

In the fibroblast subpopulations, the PRG4+, synovium-specific fibroblast population resembling synovial lining fibroblasts, showed decreased proportions in RA patients (Figure 5A) as previously described (24). However, statistically this change did not reach significance (p=0.0874). SPARC+ were present in skin and synovium with a small proportion (<2%) in lung (Figure 5B). No disease specific alteration in proportions were detected in this subtype nor in the lung-specific A2M+ fibroblast population (Figure 5C). Similarly to the SPARC+ population, the ‘pro-inflammatory’ APOE+, *CXCL12* producing fibroblasts population was high in skin and synovium and also found in lungs, but at <10% (Figure 5D). Its proportion increased in psoriatic skin compared to healthy and tended to increase in RA synovium. In contrast, this population was downregulated in fibrotic lung conditions (Figure 5D). ACTA2+ fibroblasts were mainly present in lung with a small population in heart tissue (Figure 5E). ACTA2+ fibroblasts proportions strongly increased in heart failure as well as in IPF and SSc lung involvement. Like the SPARC+ and the APOE+ populations, the MFAP5+ fibroblast population was present in skin, synovium and lung, but overall, rather decreased in disease (Figure 5F). A statistically significant decrease was seen in lung disease, as well as in acne. CDH19+ were the predominant fibroblast population in heart, but also found in lungs, skin and a small population in synovium (<10%)(Figure 5G). The CDH19+ fibroblast population decreased in heart failure, lung disease, as well as psoriasis. In contrast, the newly found CALD1+ population, which was present in all analysed tissues, but most prominent in lung and skin, increased in heart failure (Figure 5H).

**Figure 5.**
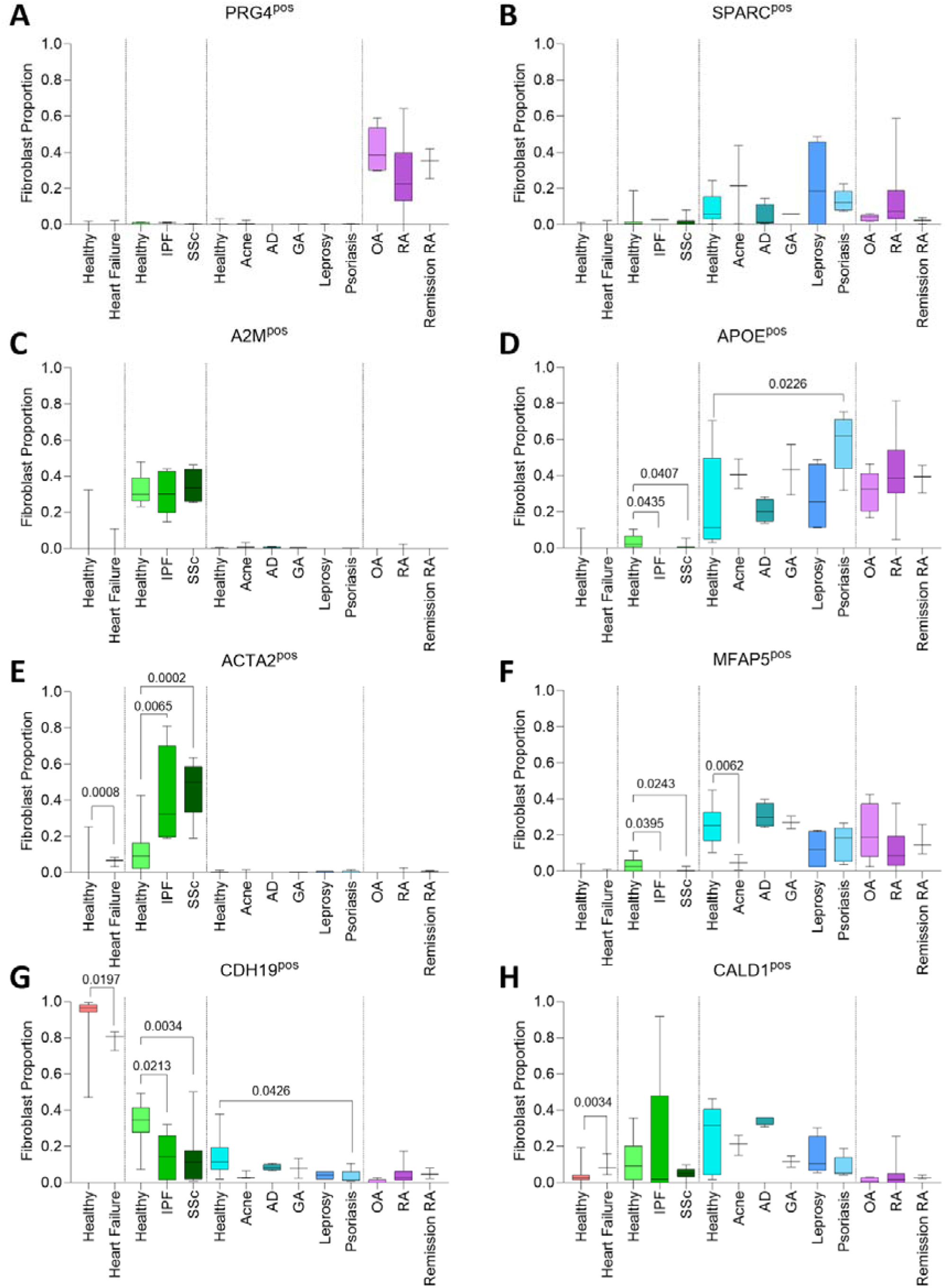
Distinct stromal cell clusters dominate different tissues and disease states. Box and whisker plots comparing the relative proportion of cells in each myeloid cluster across different tissues healthy donors and disease states. Statistical analysis applied ANOVA with Dunnett test for multiple comparison to compare healthy with disease state within each tissue. For heart t test was applied. Significant P values (*P* < 0.05) are shown. Heart failure (n=6), idiopathic pulmonary fibrosis (IPF; n=5), systemic sclerosis-associated interstitial lung disease (SSc; n=12), acne (n=4), leprosy (n=4), psoriasis (n=5), granuloma annulare (GA; n=2), atopic dermatitis (AD; n=4), osteoarthritis (OA; n=5), rheumatoid arthritis (RA; n=35) and RA in sustained clinical remission (n=5). Red = heart; green = lung; blue = skin; purple = synovium.

Even though, large sample variation between patients made comparison across disease states more difficult, we could identify common and disease specific changes of myeloid and stromal cell populations. In particular, the proportion of SPP1+ macrophages increased in diseased states in heart, lung, skin and synovium. CDH19+ and MFAP5+ fibroblast populations generally decreased in diseased states across tissues, while increased presence of APOE+ and ACTA+ fibroblast was more disease and tissue specific.

### Location of specific stroma-myeloid cell populations in diseased tissue

We then set out to identify the specific locations of the dominant, common fibroblast populations within lung, skin and heart and to visualize the identified cell-cell interactions. To identify SPARC+ fibroblasts in the tissue, we used periostin (POSTN), which appeared as prominent cell-specific marker gene for this population. To determine the presence of APOE+ fibroblasts within tissues, we stained for CXCL12. CXCL12, also called stromal cell-derived factor 1, is highly expressed by APOE+ fibroblasts and is the major chemokine secreted by fibroblasts. Finally, we stained Microfibrillar-associated protein 5 (MFAP5) to detect the MFAP5+ subpopulation.

As suggested by the transcriptome analysis, MFAP5 and periostin expression was highest in skin and synovium (Figure 7A) whereas periostin expression was low in heart and lung tissue (Supplemental Figure 4). In healthy as well as ischemic heart tissue, MFAP5 formed a thin layer around cardiac muscle cells. In skin and synovium, periostin and MFAP5 expression was confined to distinct and remotely exclusive compartments (Figure 7). In the skin, periostin was localized in the papillary dermis, while MFAP5 was expressed in the reticular dermis. In the synovium, periostin expression was found mainly in close association to blood vessels, while MFAP5 was detected throughout the remaining sublining area. This distinct pattern of distribution was maintained in psoriatic skin and RA synovial tissue. Thus, expression of these marker proteins supported the presence of two distinct fibroblast subpopulations, creating specialized tissue niches in skin as well as synovium in health and disease.

**Figure 7.**
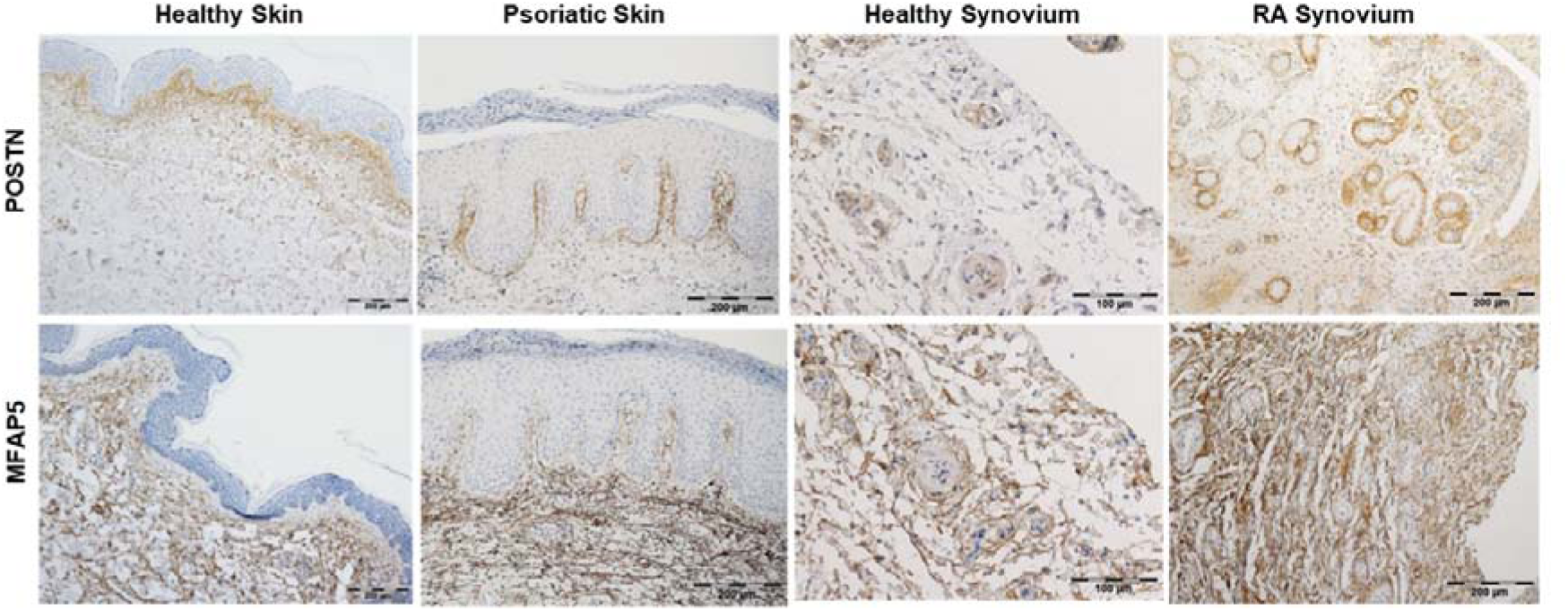
Expression of POSTN, MFAP5, CXCL12 and SPP1 in skin and synovium. Immunohistochemical staining of POSTN and MFAP5 in healthy skin, psoriatic skin, healthy synovium and RA synovium. Magnification 100x for skin and RA synovium, 200x for healthy synovium.

## Discussion

In this work, we created a myeloid and stromal single cell atlas for the dissection of tissue macrophage and fibroblast phenotype across heart, lung, skin, and synovium. We identified tissue unique and shared clusters of myeloid cells and fibroblasts and could demonstrate disease specific changes of these subpopulations.

We tested three distinct approaches for cross-tissue integration of publicly available scRNAseq data and identified Harmony as the standout method. Surprisingly, the BBKNN method, which is the common choice for integration of large scale scRNAseq datasets (29, 30), was the least effective when applied during preliminary testing. Furthermore, a recent benchmarking study assessed the performance of different tools in five different integration scenarios and identified Harmony and Seurat V3 as the top methods (31). This emphasizes that selection of methods for integration is dependent on dataset characteristics (e.g., cell number, diversity) and thus highlights the benefits of testing more than one approach.

Combined data analysis can be challenging as cellular gene expression profiles are altered by differences of, not only important biological (donor, tissue, disease) factors, but also confounding technical (protocol, sequencing platform, location) variables. However, we are confident that our data integration was effective as we see populations previously known to be unique to tissues. Langerhans cells were the only tissue unique myeloid cell population, whereas we could identify three tissue unique fibroblast cell populations, which presumably fulfil tissue specialized functions. PRG4+ fibroblasts have previously been described as the synovial fibroblast population that form the intimal, lining layer of the synovium (24). In accordance with their function to produce the lubricants for the synovial fluid, they are characterized by high levels of hyaluronan synthase 1 (*HAS1*). In health, ACTA2+ fibroblasts were present exclusively in lung tissue, as were A2M+ fibroblasts. Fibroblasts producing high levels of *ACTA2* and their expansion in lung fibrosis were previously described in several scRNAseq studies of lung tissues (7, 32, 33). The A2M+ lung fibroblasts in our dataset most likely correspond to a previously identified fibroblasts subtype in the lung found near alveoli (33), both strongly expressing *GPC3*. Their specific function is however unknown. Strikingly there was a strong enrichment of CDH19+ fibroblasts in the heart. While we cannot exclude that this enrichment is due to technical limitations, it is noteworthy that previously cardiac fibroblasts expressing high levels of *C7* and *ADH1D*, like our CDH19+ fibroblasts, were described as one of the main fibroblast subtypes found in cardiac ventricles (13). Furthermore, in the human protein atlas, *CDH19* is annotated in the ‘cardiac muscle contraction’ cluster and was assigned to fibroblasts (in addition to oligodendrocytes and melanocytes). Thus, this cluster might represent a specific cardiac fibroblast subtype with a specialised role in supporting the contracting myocardium.

Shared fibroblast populations were most evident in skin and synovium (SPARC+, APOE+, MFAP5+). Fibroblasts subpopulations with similar marker genes as our SPARC+ and MFAP5+ fibroblasts were also described in a cross-tissue analysis of gut, lung, salivary gland and synovium (34) and might thus represent universal archetypes of fibroblasts. Our staining of MFAP5 and POSTN in the skin and synovium corroborate the concept that these two presumably functionally distinct fibroblast subpopulations exist across tissues and form anatomically defined niches. Functional differences between papillary and reticular skin fibroblasts were previously described (35). Our reticular MFAP5+ fibroblast corresponds to a skin fibroblast subpopulation that has been connected to skin homeostasis and wound healing (18, 20, 36). Whether this fibroblast population plays the same role in other tissues remains to be examined. However, our data showing that the proportion of this fibroblast subtype tends to be reduced in disease in the analysed tissues suggest that a proportional loss of this population may be of pathogenic relevance.

In summary, we successfully integrated 14 different datasets generated from four different tissues and spanning several diseases. We show that fibroblast subpopulations in these tissues are more heterogeneous than myeloid cells. Nevertheless, skin and synovium in particular share several fibroblast populations. Since these are impacted by chronic inflammation, they could be of interest for further functional studies, especially in diseases that occur in both organs, such as psoriatic arthritis.

## Methods

### Quality checks, filtering, and clustering of lung, skin, heart and synovial tissue scRNAseq datasets

We collected scRNAseq data from 14 public datasets shown in Figure 1A, spanning 4 distinct tissues (heart, lung, skin, and synovium) including 162 male and female healthy donors and disease patients, with conditions such as – Heart Failure (HF) (15), Idiopathic Pulmonary Fibrosis (IPF) (7), Systemic Sclerosis-associated Interstitial Lung Disease (SSc-ILD) (7, 17), Acne (19), Leprosy (19), Psoriasis (19), Granuloma Annulare (GA)(19), Atopic Dermatitis (AD) (18), Osteoarthritis (OA) (24), Rheumatoid Arthritis (RA) (21–24) and RA that is in sustained clinical remission (21). We began by creating an atlas for each tissue, combining individual datasets Seurat’s anchor integration strategy (25). Read count matrices were obtained via the links in Figure 1A and were read into R. Prior to Seurat object creation, we queried and converted gene annotation to Ensembl transcripts (ENST) to using the gprofiler2 package (gconvert) to ensure all datasets had the same gene annotation. We then followed the standard protocol for pre-processing and clustering of scRNAseq data using the Seurat package (3.1.2) in R. UMAP based on PCA cell embeddings were generated for each dataset and the first 30 principal components (PCs) were used for visualization (RunUMAP). The same PCs were used in determination of the k-nearest neighbors for each cell during SNN graph construction before clustering at a chosen resolution of 0.2 (FindNeighbors, FindClusters). By investigation of canonical marker genes, we identified macro clusters of myeloid (*MARCO, CD14, LYZ*), lymphocytes (*CD3D*) and stromal (*COL1A1, PDPN*) cells and isolated these populations for downstream integration. We then performed integration of datasets from the same tissue using Seurat’s anchor integration strategy. We integrated all genes that are common between datasets, using the functions: FindIntegrationAnchors, and IntegrateData (features.to.intergrate to find all common genes). These “integrated” batch-corrected values were then set as the default assay and the gene expression values are scaled before running principle component analysis. UMAP was computed based on PCA cell embeddings was generated from integrated counts batch-normalized by Seurat and the first 30 principle components (PCs) are visualized (RunUMAP). The same PCs were used in determination of the k-nearest neighbours for each cell during SNN graph construction before clustering at a chosen resolution of 0.4 (FindNeighbors, FindClusters). The result of such visualization and the total number of cells in each tissue atlas is shown in Figure 1B.

### Integration of scRNAseq data of lung, skin, heart and synovial tissue myeloid cells and fibroblasts

To optimize our pipeline for cross-tissue integration, we tested the ability of three distinct methods (Seurat (25), Harmony (26) and BBKNN (27) to integrate a random subset myeloid (n=4500), stromal (n=4500) and T cells (n=1000) cells from each tissue.

*Subsampling.* For reproducibility, we set a seed value of 111. For each tissue, we identified the name of the cells in each lineage (WhichCells) and specified the number of cell names we wished to subsample (using the base R sample function) before isolating the reduced number of myeloid, stromal and T cells using Seurat’s subset function. After subsampling, we merged the subsampled (10,000 cells) macrophage-fibroblast-tcell (MFT) Seurat object from each tissue to visualize how the data looks pre-integration. We did so by computing the UMAP based on PCA cell embeddings generated from non-batch-normalized counts and visualizing the first 30 PCs (RunUMAP).

*Comparison of Integration Methods*. Firstly, we applied Seurat’s anchor integration strategy4 using the same process described above for the integration of datasets from the same tissue. After computing UMAP visualization based onSeurat batch-normalized counts, we compared how cells from distinct tissues and lineages overlapped (DimPlot). Next, we applied the Harmony5 (0.1.0) protocol to integrate by sample (RunHarmony). This function creates a batch corrected PCA which is then used for downstream visualization and clustering (RunUMAP, FindNeighbors, FindClusters). Finally, we applied BBKNN6 (1.3.6), an integration method that is implanted in python. To convert our subsampled MFT Seurat object for use in python, we used the SeuratDisk (0.0.0.9019) package. This process involved saving our Seurat object as .h5Seurat file (SaveH5Seurat) before converting to .h5ad file (Convert). The .h5ad file can then be read into python using the scanpy7 (1.5.0) package, which creates an AnnData object (sc.read). Before running BBKNN, we used scanpy to create PCA (sc.tl.pca, sc.pl.pca) and UMAP visualization (sc.tl.umap, sc.pl.umap). We then ran BBKNN to integrate by sample (sc.external.pp.bbknn) before re-computing UMAP visualization (sc.tl.umap, sc.pl.umap). To continue with downstream clustering and plotting, we converted our AnnData object in python back into Seurat object in R. The AnnData object was saved in python as .h5ad file (anndata.write) before using SeuratDisk to convert to .h5Seurat file (Convert) which could be loaded into R and saved as .rds (LoadH5Seurat, saveRDS). We then used the Seurat package for clustering (FindNeighbors, FindClusters).

*Harmony integration of all cells.* After interrogating the output of distinct integration methods, we decided to proceed with Harmony for further analysis. We repeated the integration protocol as before, this time using all myeloid, stromal and T cells from each tissue. After integrating by sample (RunHarmony), we used the top 50 PC of the resultant harmony batch corrected PCA for UMAP visualization and a resolution of 0.8 for clustering. Clusters were identified by interrogation of marker genes and T lymphocyte and any contaminant or doublet clusters were removed, leaving a total of 253,843 macrophage and fibroblast cells across tissues. The remaining clusters were renamed.

*Differential Expression Analysis*. To identify cluster markers, the Seurat function FindAllMarkers was used with “test.use=MAST”. As recommended in the best practice of Seurat, for DE comparison the non-batch normalized counts were used. For identification of cluster markers, we specify that any markers identified must be expressed by at least 40% of cells in the cluster (‘min.pct’ parameter 0.4). We use the default values for all other parameters. Genes are considered significantly DE if the adjusted p-value (< 0.05) by Bonferroni Correction and multiple test correction (multiple by number of tests). To visualise heatmaps the pheatmap package was adapted.

*Statistical evaluation of cluster proportions across tissues*. Statistical analyses were performed between healthy state of each tissue, using one-way ANOVA with Kruskal-Wallis test for multiple comparisons and significant (P < 0.05) comparisons plotted.

### Immunhistochchemical analysis of heart, lung, skin and synovial tissue

Skin, synovium, heart and lung were fixed in 10% formalin and embedded in paraffin. Immunohistochemical single staining for periostin and MFAP5 was performed using the BOND-Max Fully Automated Immunohistochemistry system (Leica Biosystems). After deparaffinisation, tissues were pretreated with the Epitope Retrieval Solution 2 (EDTA-buffer pH 9) at 100°C for 30 min. Slides were then washed, and peroxidase blocking solution was applied. Slides were again washed and incubated with the primary antibody (polyclonal periostin human anti-rabbit, Abcam ab14041, dilution 1:2000 in Bond Primary Antibody Diluent or polyclonal MFAP5 human anti-rabbit, Invitrogen PA5-52706, dilution 1:2000 in Bond Primary Antibody Diluent) for 30min. Washing steps were repeated and slides were incubated with polymer poly-HRP anti-rabbit IgG antibody for 10 min. Colour development was carried out with DAB (diaminobenzidine) reagent for 10 min. Finally, tissues were counterstained with hematoxylin for 10 min and washed with wash buffer and DI water.

For the double immunohistochemical staining with CXCL12 and osteopontin in synovium and skin sections, the sequential double staining protocol of BOND-Max Fully Automated Immunohistochemistry system (Leica Biosystems) was used. After deparaffinisation, tissues were pretreated with the Epitope Retrieval Solution 1 (citrate-buffer pH 6) at 100°C for 30 min. Slides were then washed, and incubated with the first primary antibody (polyclonal human anti-rabbit SPP1 antibody, Abcam ab8448, dilution 1:1000 in Bond Primary Antibody Diluent) for 30 min, followed by the polymerisation step with polymer poly-AP anti-rabbit IgG antibody for 10 min. Colour development was carried out with Fast Red substrate chromogen for 10 min. Subsequently, peroxidase blocking solution was added, followed by incubation with the second primary antibody (human anti-mouse CXCL12 antibody, R&D Systems clone 79018, dilution 1:400 in Bond Primary Antibody Diluent) for 30 min. An additional incubation step with a rabbit anti-mouse post-primary antibody for 30 min was required. Slides were washed and incubated with polymer poly-HRP anti-rabbit IgG antibody for 10 min. Colour development was carried out with DAB (diaminobenzidine) reagent for 10 min. Finally, tissues were counterstained with hematoxylin for 10 min and washed with wash buffer and DI water. Images were acquired using the Olympus BX53 microscope.

## Author contributions

LMD: acquiring data, analysing data, writing the manuscript; OH: analysing data; MT: conducting experiments; GK: designing research studies, writing the manuscript; PB: designing research studies, writing the manuscript; RM: acquiring data, analysing data, writing the manuscript; TDO: designing research studies, writing the manuscript; MKS: designing research studies, writing the manuscript; CO: designing research studies, writing the manuscript

