## Supplemental Figure for "Human Fibroblast-Myeloid cell tissue atlas across lung, synovium, skin and heart"

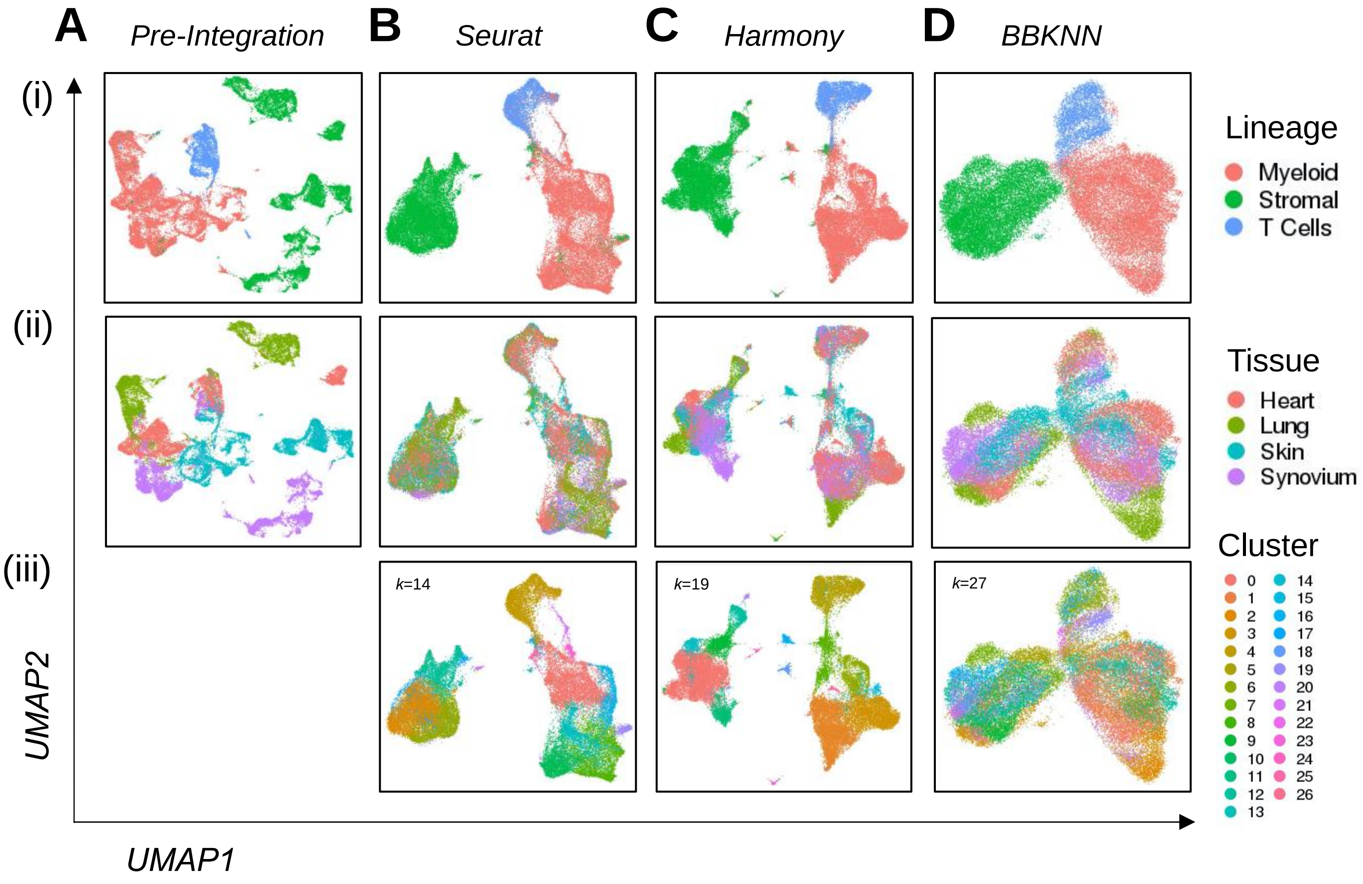


**Supplemental Figure 1. Optimizing the normalized integration strategy of myeloid (4.5k), stromal (4.5k) and T cells (1k) from heart, lung, skin and synovial tissue scRNAseq data.** UMAP visualization of combined analysis of scRNAseq data from distinct tissues with cells coloured by (i) lineage, (ii) tissue of origin, and (iii) cluster identity. Comparison of UMAP visualization is shown for 40,000 cells combined from heart, lung, skin and synovium with a normalized number of myeloid (n=4500), stromal (n=4500) and T cells (n=1000) cells randomly subsampled from each tissue. UMAP visualization is compared (A) pre-integration and post-integration with (B) Seurat, (C) Harmony, and (D) BBKNN.

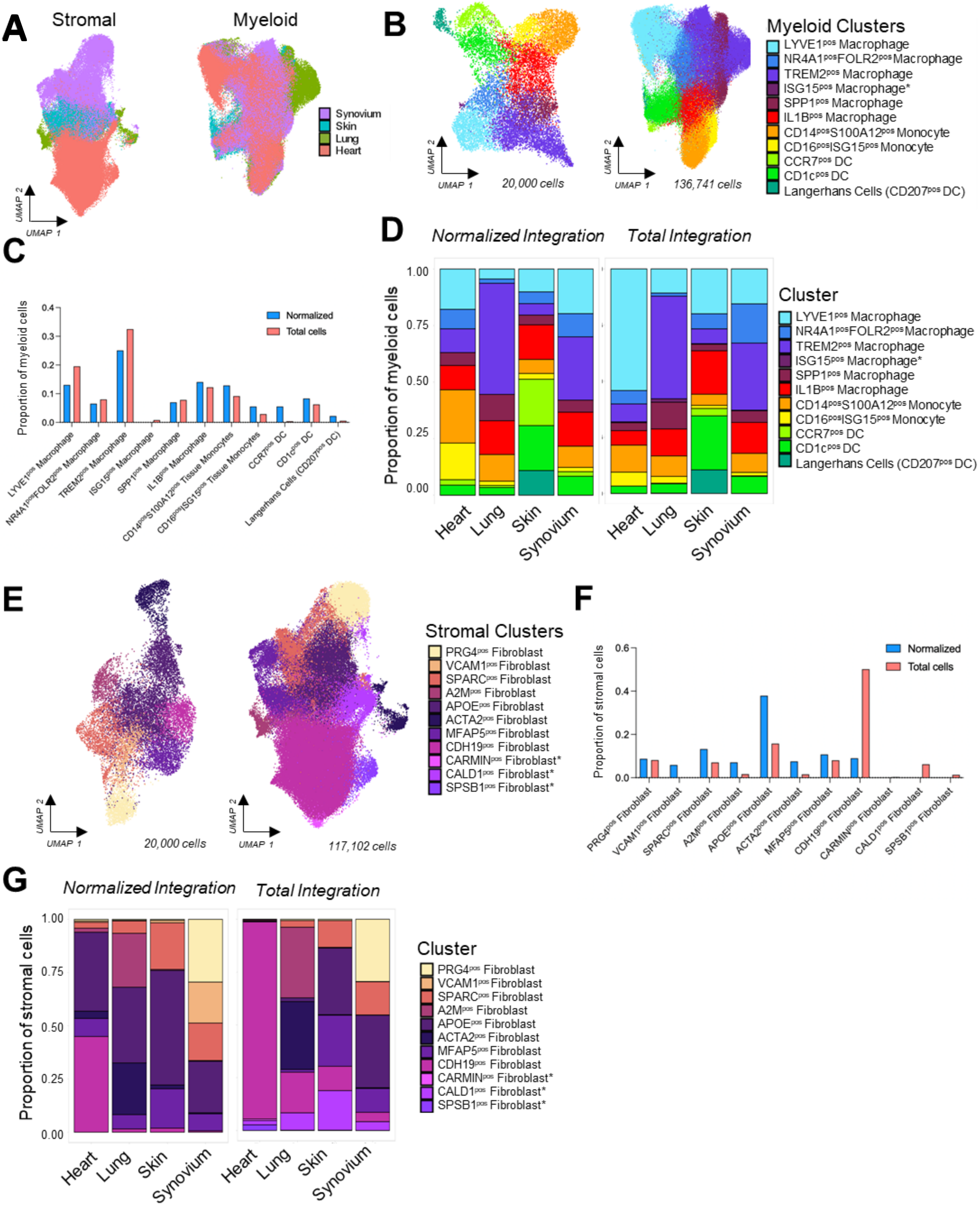


**Supplemental Figure 2. Harmony integration of all myeloid and stromal cells from each tissue to facilitate cell cluster proportion analysis. (A)** UMAP visualization of 136,741 myeloid and 117,102 stromal cells from cross-tissue integration with Harmony. Cells are represented by individual dots and coloured by tissue origin. **(B)** Comparison of UMAP visualization of myeloid cells from normalized and total integration. Cells coloured by cluster identity. **(C)** Bar plot comparing the relative proportion of each cluster of all myeloid cells in both normalized and total tissue atlas. **(D)** Stacked bar plot of relative proportion of cells in each myeloid cluster across different tissue in normalized and total integration. **(E)** Comparison of UMAP visualization of stromal cells from normalized and total integration. Cells coloured by cluster identity. **(F)** Bar plot comparing the relative proportion of each cluster of all stromal cells in both normalized and total tissue atlas. **(G)** Stacked bar plot of relative proportion of cells in each stromal cluster across different tissue in normalized and total integration. **(H)** Dendrogram visualization of hierarchical clustering of stromal cell populations

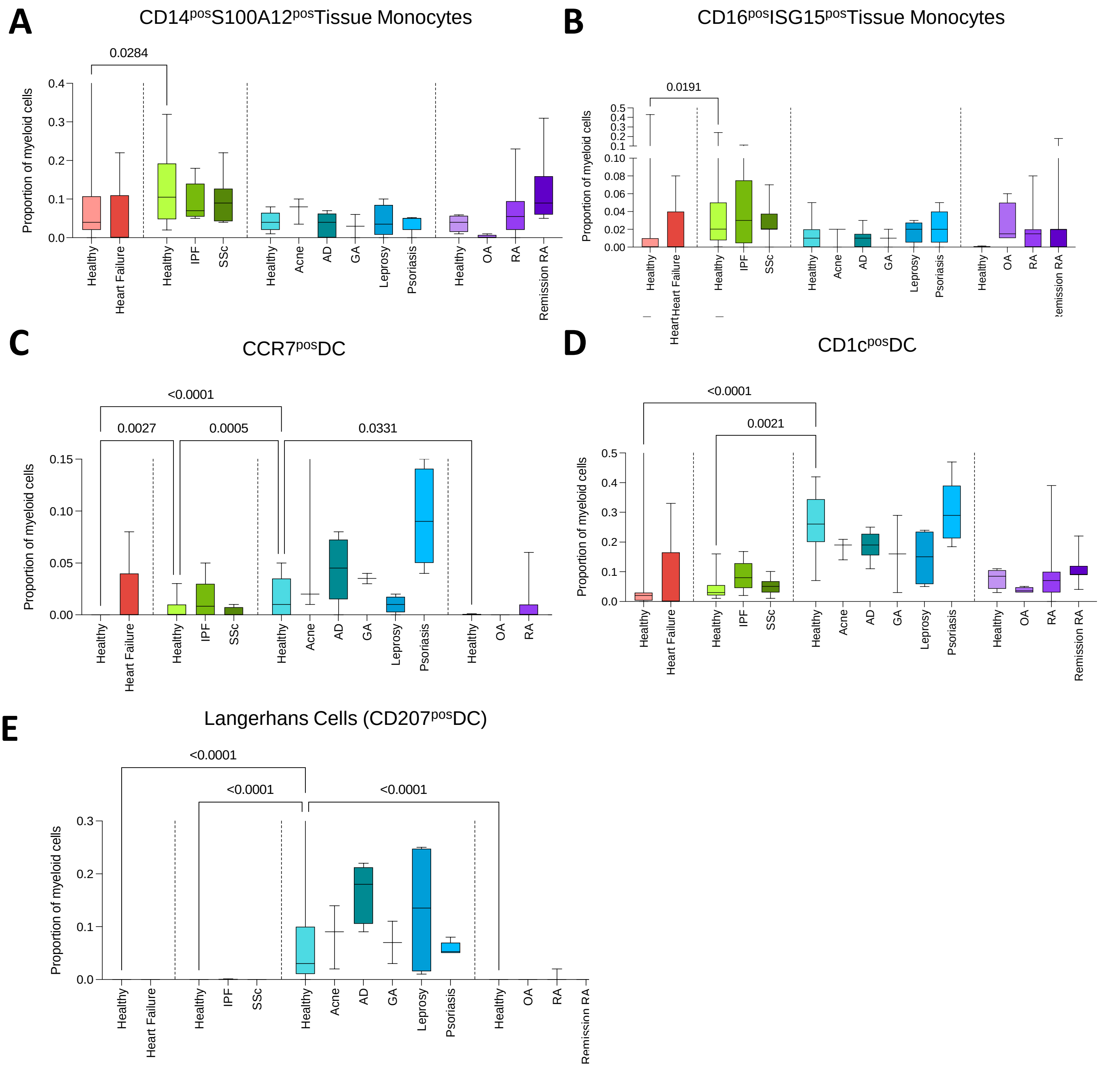


**Supplemental Figure 3. Distinct monocyte and dendritic cell clusters in different tissues and disease states.** Box and whisker plots comparing the relative proportion of cells in each myeloid cluster across different tissues healthy donors and disease states. Statistical analysis applied ANOVA with Kruskal-Wallis test for multiple comparison to compare healthy state of each tissue. Significant P values ($P < 0.05$) are shown. Heart failure (n=6), idiopathic pulmonary fibrosis (IPF; n=5), systemic sclerosis-associated interstitial lung disease (SSc; n=12), acne (n=4), leprosy (n=4), psoriasis (n=5), granuloma annulare (GA; n=2), atopic dermatitis (AD; n=4), osteoarthritis (OA; n=5), rheumatoid arthritis (RA; n=35) and RA in sustained clinical remission (n=5). Red = heart; green = lung; blue = skin; purple = synovium

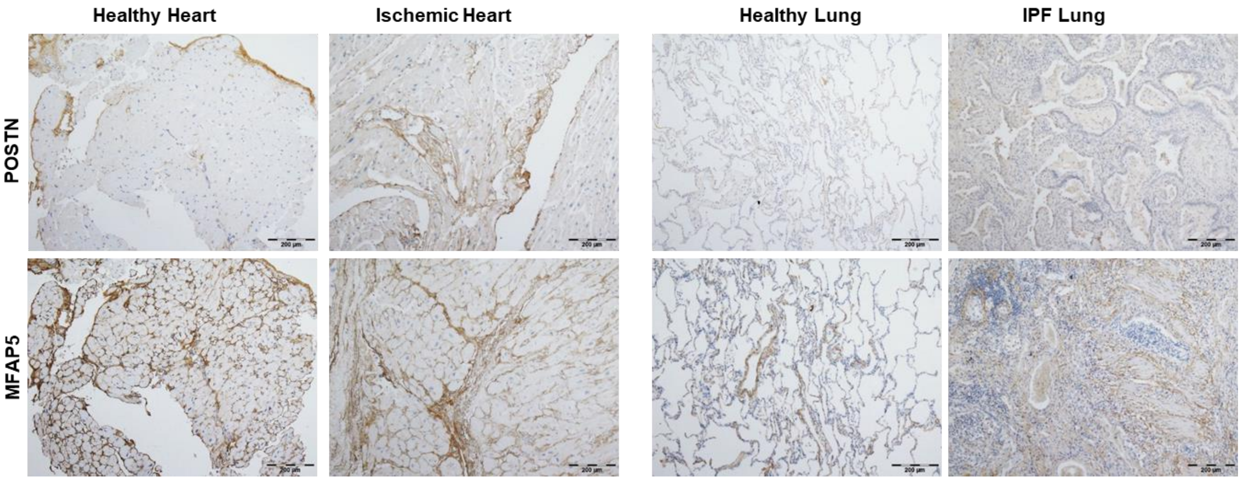


**Supplemental Figure 4. Expression of POSTN and MFAP5 in heart and lung tissue.** Immunohistochemical staining of POSTN and MFAP5 in healthy heart, ischemic heart, healthy lung and ischemic lung. Magnification 100x.
